# Computational analysis of microRNA-mediated interactions in SARS-CoV-2 infection

**DOI:** 10.1101/2020.03.15.992438

**Authors:** Müşerref Duygu Saçar Demirci, Aysun Adan

**Affiliations:** Bioinformatics, Abdullah Gül University, Kayseri, Turkey; Molecular Biology and Genetics, Abdullah Gül University, Kayseri, Turkey

## Abstract

MicroRNAs (miRNAs) are post-transcriptional regulators of gene expression that have been found in more than 200 diverse organisms. Although it is still not fully established if RNA viruses could generate miRNAs that would target their own genes or alter the host gene expression, there are examples of miRNAs functioning as an antiviral defense mechanism. In the case of Severe acute respiratory syndrome coronavirus 2 (SARS-CoV-2), there are several mechanisms that would make miRNAs impact the virus, like interfering with replication, translation and even modulating the host expression. In this study, we performed a machine learning based miRNA prediction analysis for the SARS-CoV-2 genome to identify miRNA-like hairpins and searched for potential miRNA – based interactions between the viral miRNAs and human genes and human miRNAs and viral genes. Our PANTHER gene function analysis results indicate that viral derived miRNA candidates could target various human genes involved in crucial cellular processes including transcription. For instance, a transcriptional regulator, STAT1 and transcription machinery might be targeted by virus-derived miRNAs. In addition, many known human miRNAs appear to be able to target viral genes. Considering the fact that miRNA-based therapies have been successful before, comprehending mode of actions of miRNAs and their possible roles during SARS-CoV-2 infections could create new opportunities for the development and improvement of new therapeutics.

## Introduction

Coronoviruses (CoVs) are pathogens with serious health effects including enteric, respiratory, hepatic and central nervous diseases on human and animals. Zoonotic CoVs, Severe Acute Respiratory Syndrome-Coronavirus (SARS-CoV) and Middle East Respiratory Syndrome coronavirus (MERS-CoV), have been identified the sources of outbreaks in 2002/2003 and 2012, respectively with high mortality rates due to severe respiratory syndrome ^1,2^. Besides zoonotic CoVs, there are four types of human CoVs have been identified known as HCoV-OC43, HCoV-2293, HCoV-NL63 and HCoV-HKU1 ^1^. Unknown pneumonia have been detected in Wuhan, China and spread globally since December 2019. The World Health Organization (WHO) named this new coronavirus as Severe Acute Respiratory Syndrome Coronavirus 2 (SARS-CoV-2) responsible for the new disease termed Coronavirus Disease 2019 (COVID-19) and it is the seventh identified CoV with animal origin infecting human ^2^.

Coronaviruses are positive-single stranded RNA (+ssRNA) viruses with exceptionally large genomes of ∼ 30 kb with 5′cap structure and 3′polyA tail. The coronavirus subfamily is divided into four genera: α, β, γ, and δ based on serotype and genome features ^3^. The genome of a typical CoV codes for at least 6 different open reading frames (ORFs), which has variations based on the CoV type ^4^. Some ORFs encode non-structural proteins while others code for structural proteins required for viral replication and pathogenesis. Structural proteins include spike (S) glycoprotein, matrix (M) protein, small envelope (E) protein, and nucleocapsid (N) protein with various roles for virus enterance and spread. SARS-CoV-2 belongs to β CoV with 45-90% genetic similarity to SARS-CoV based on sequence analysis and might share similar viral genomic and transcriptomic complexity ^3,5^. Currently, it has been also revealed that SARS-COV-2 has a very high homology with bat CoVs, which indicated how it is transmitted to human without knowing intermediate carriers ^6^. S protein of SARS-CoV-2 has a strong interaction with human angiotensin-converting enzyme 2 (ACE2) expressed on alveolar epithelial cells which shows the way of virus infection in human ^7^.

MicroRNAs (miRNAs) are small, noncoding RNAs that play role in regulation of the gene expression in various organisms ranging from viruses to higher eukaryotes. It has been estimated that miRNAs might influence around 60% of mammalian genes and their main effect is on regulatory pathways including cancer, apoptosis, metabolism and development ^8^.

Although the current release of miRNAs, the standard miRNA depository, lists miRNAs of 271 organisms, only 34 of them are viruses ^9^. While, the first virus-encoded miRNAs was discovered for the human Epstein-Barr virus (EBV) ^10^, more than 320 viral miRNA precursors were reported so far. Although it has been shown that various DNA viruses express miRNAs, it is still debatable if RNA viruses could also encode. The major concerns regarding miRNAs of RNA viruses are based on ^11^:

- the fact that RNA viruses that replicate in cytoplasm, do not have access to nuclear miRNA machinery
- since RNA is the genetic material, miRNA production would interfere with viral replication

On the other hand, many features of miRNA - based gene regulation seems to be especially beneficial for viruses. For instance, through targeting specific human genes by viral miRNAs, it is possible to form an environment suitable for survival and replication of the virus. Furthermore, for viral miRNAs it is likely to escape host defense system since host itself generates miRNAs in the same manner.

Currently, options for the prevention and treatment of CoVs are very limited due to the complexity. Therefore, detailed analysis of CoV-host interactions is quite important to understand viral pathogenesis and to determine the outcomes of infection. Although there are studies regarding to the viral replication and their interaction with host innate immune system, the role of miRNA-mediated RNA-silencing in SARS-CoV-2 infection has not been enlightened yet. In this study, SARS-CoV-2 genome was searched for miRNA-like sequences and potential host-virus interactions based on miRNA actions were analyzed.

### Methods

Data analysis, pre-miRNA prediction, mature miRNA detection workflows were generated by using the Konstanz Information Miner (KNIME) platform ^12^

### Data

Genome data of the virus was obtained from NCBI:

Severe acute respiratory syndrome coronavirus 2 isolate Wuhan-Hu-1, complete genome

GenBank: MN908947.3

MiRNA prediction workflow izMiR ^13^ and its related data were taken from Mendeley Data: https://data.mendeley.com/datasets/mgh5r9wny7/1

Mature miRNA sequences of human were downloaded from miRBase Release 22.1 ^14^.

### Pre-miRNA prediction

Genome sequences of SARS-CoV-2 were transcribed (T->U) and divided into 500 nt long fragments with 250 overlaps. Then these fragments were folded into their secondary structures by using RNAfold ^15^ with default settings and hairpin structures were extracted, producing 950 hairpins in total.

A modified version of izMiR (SVM classifier is changed to Random Forest and latest miRBase version was used for learning) was applied to these hairpins with ranging lengths (from 7 to 176) (Figure 1). Based on the mean value of averages of 3 classifiers’ prediction scores (Decision Tree, Naive Bayes and Random Forest), 29 hairpins passed 0.900 threshold and used for further analysis.

**Figure 1:**
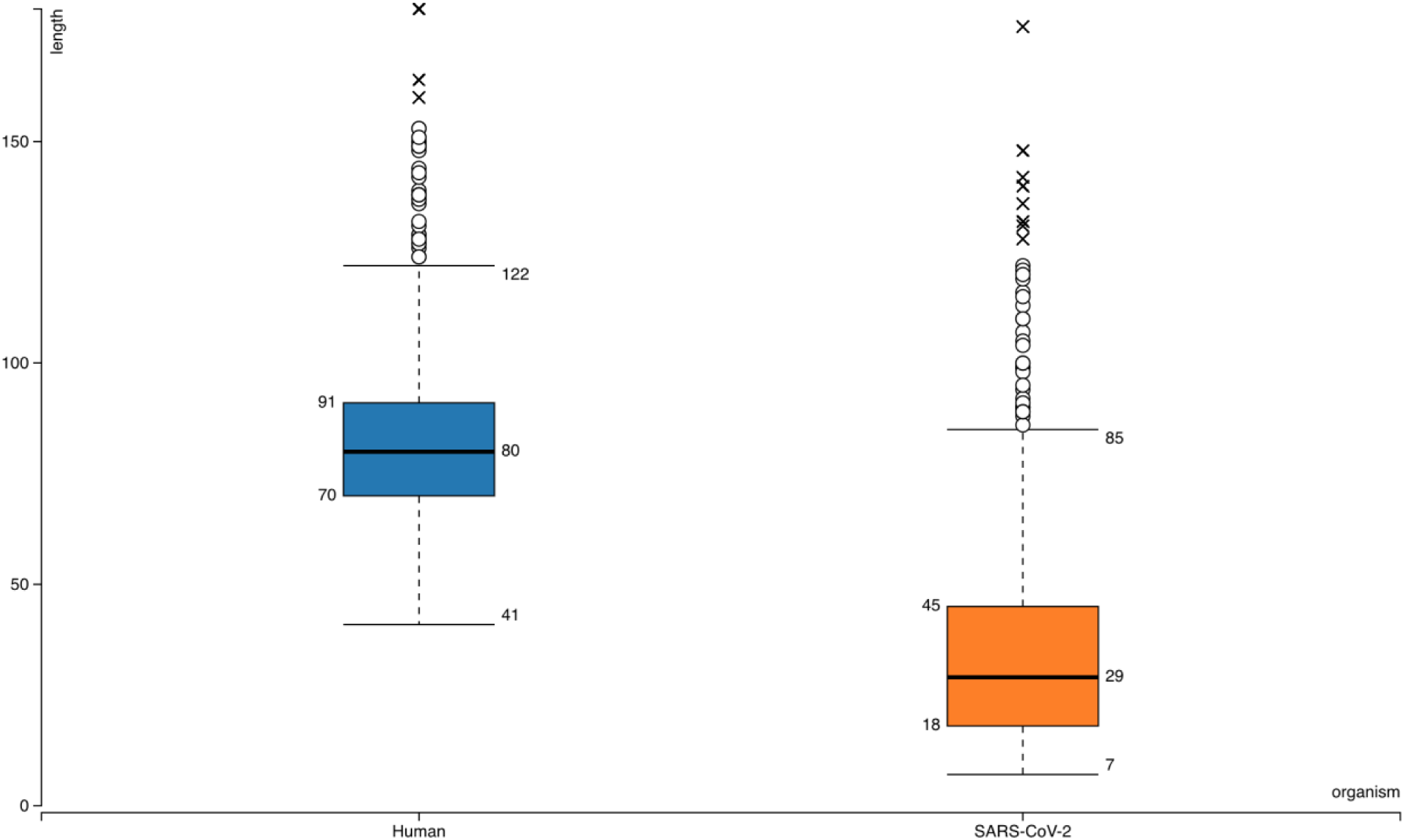
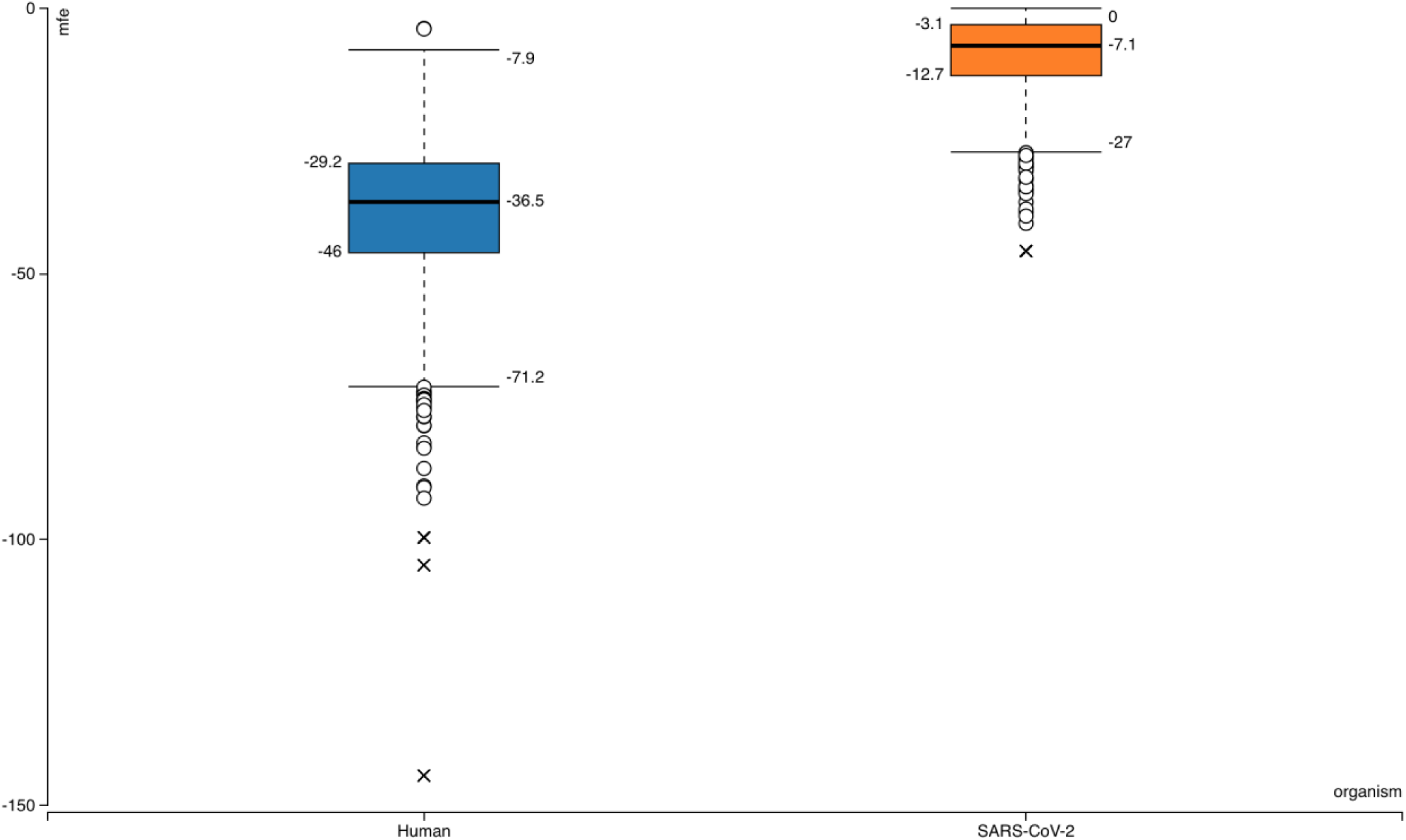
Box-plots for comparison of general features of human pre-miRNAs and SARS-CoV-2 hairpins. Hairpin lengths shown in top graph. Minimum free energy scores obtained from RNAfold ^15^ plotted in bottom panel.

### Mature miRNA prediction and targeting

Selected hairpins were further processed into smaller sequences; maximum 23 nt length with 6 nt overlaps. Then, these fragments were filtered based on minimum length of 15 and their location on the hairpins (sequences not involving any loop nucleotides were included). Target search of these remaining 30 candidate mature miRNAs were performed against human and SARS-CoV-2 genes by using psRNATarget tool with default settings ^16^. Moreover, human mature miRNAs’ from miRBase were applied for searching their targets in SARS-CoV-2 genes.

### Gene Ontology

The targets of viral miRNAs in human genes were further analyzed for their Gene Ontology (GO). To achieve this, PANTHER Classification System (http://www.pantherdb.org) was used ^17^.

## Results

Searching SARS-CoV-2 genome for sequences forming hairpin structures resulted in 950 hairpins with varying lengths (Supplementary Files). In order to use machine learning based miRNA prediction approach of izMiR workflows, hundreds of features were calculated for all of the pre-miRNA candidate sequences. Among those, minimum free energy (mfe) values required for the folding of secondary structures of hairpin sequences and hairpin sequence lengths of known human miRNAs from miRBase and predicted hairpins of SARS-CoV-2 were compared (Figure 1). Based on the box-plots shown in Figure 1, most of the extracted viral hairpins seem to be smaller than human miRNA precursors.

Since viruses would need to use at least some members of host miRNA biogenesis pathway elements, viral miRNAs should be similar to host miRNAs to a certain degree. Therefore, a classification scheme trained with known human miRNAs was applied on SARS-CoV-2 hairpins. Only 29 hairpins out of 950 passed the 0.900 prediction score threshold and used for further analysis. From these hairpins, 30 mature miRNA candidates were extracted and their possible targets for human and SARS-CoV-2 genes were investigated.

SARS-CoV-2 miRNA candidates were further analyzed to test if they were similar to any of the known mature miRNAs from 271 organism listed in miRBase. To achieve this, a basic similarity search was performed based on the Levenshtein distance calculations in KNIME. However, there was no significant similarity between hairpin or mature sequences.

While predicted mature miRNAs of SARS-CoV-2 were used to find their targets in SARS-CoV-2 and human genes, mature miRNAs of human were also applied on SARS-CoV-2 genes (Table 1, Supplementary Files). Although miRNA based self-regulation of viral gene expression is a hypothetical case, SARS-CoV-2 ORF1ab polyprotein gene might be the only one that could be a target of viral miRNAs. In total 1367 human genes seem to be targeted by viral miRNAs. Table 1 lists some of the predicted targets of SARS-CoV-2 miRNAs in human genes that have roles in transcription. The full list of miRNA – target predictions are available in Supplementary Files.

**Table 1:**
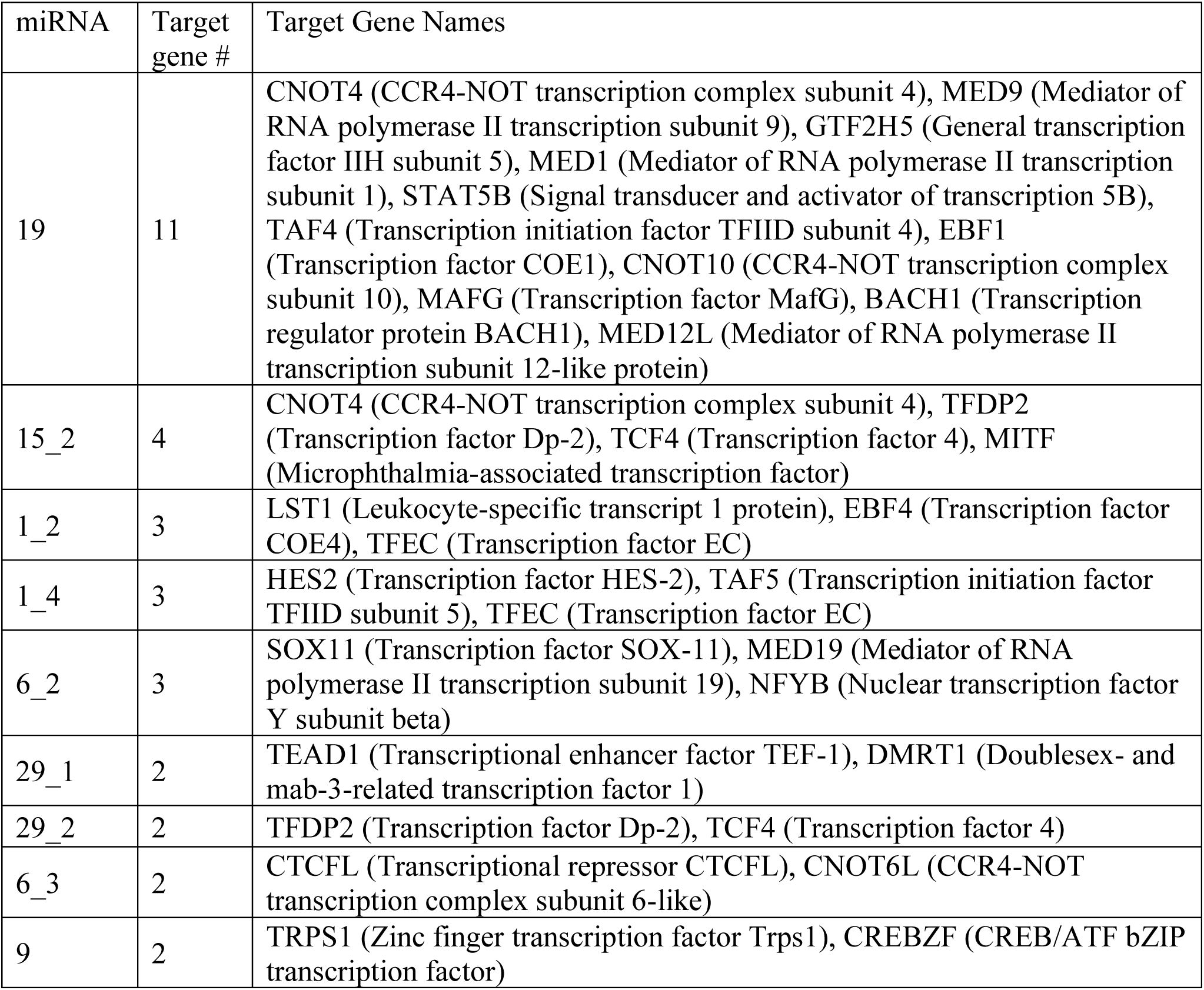
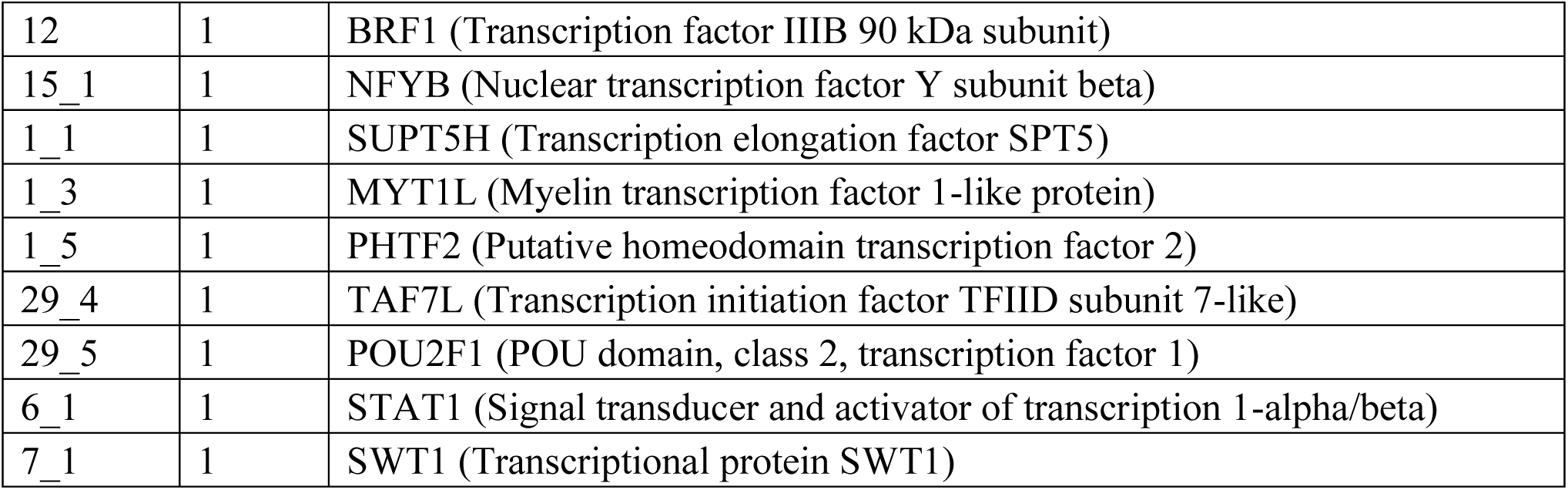
Transcription related human gene targets of viral miRNAs. MiRNA column indicates the accession numbers of mature viral miRNAs; Target gene # column shows the total number of different genes involved in transcription and targeted by the corresponding viral MiRNAs.

Considering the potential of miRNAs as biomarkers and therapeutic agents, it is essential to check if any of the known human miRNAs could target viral genes. Among 2654 mature entries of *Homo sapiens* in miRBase Release 22.1, 479 of them could target SARS-CoV-2 genes (Supplementary Files). While Envelope and ORF6 genes were targeted by single miRNAs, ORF1ab appeared to be the target of 369 different human mature miRNAs (Table 2). As expected, number of targeting events appear to be correlated with the gene length.

**Table 2:**
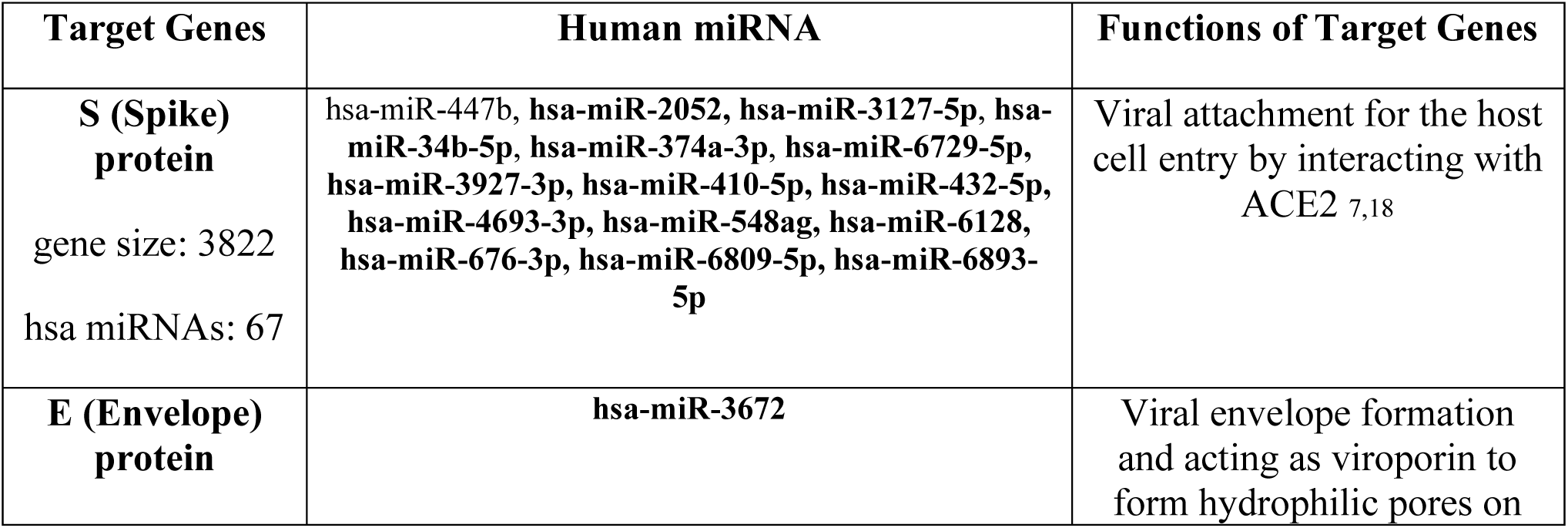
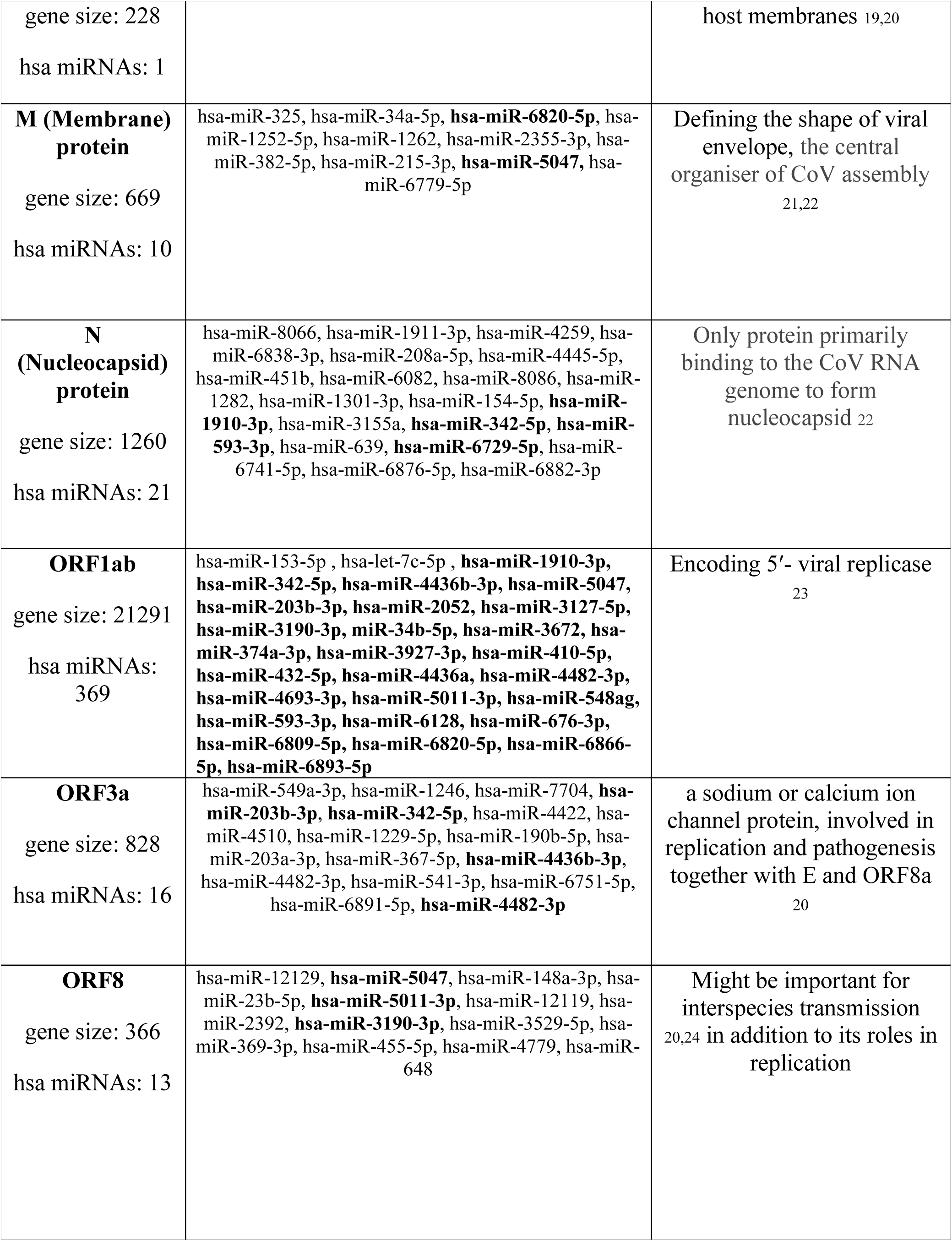
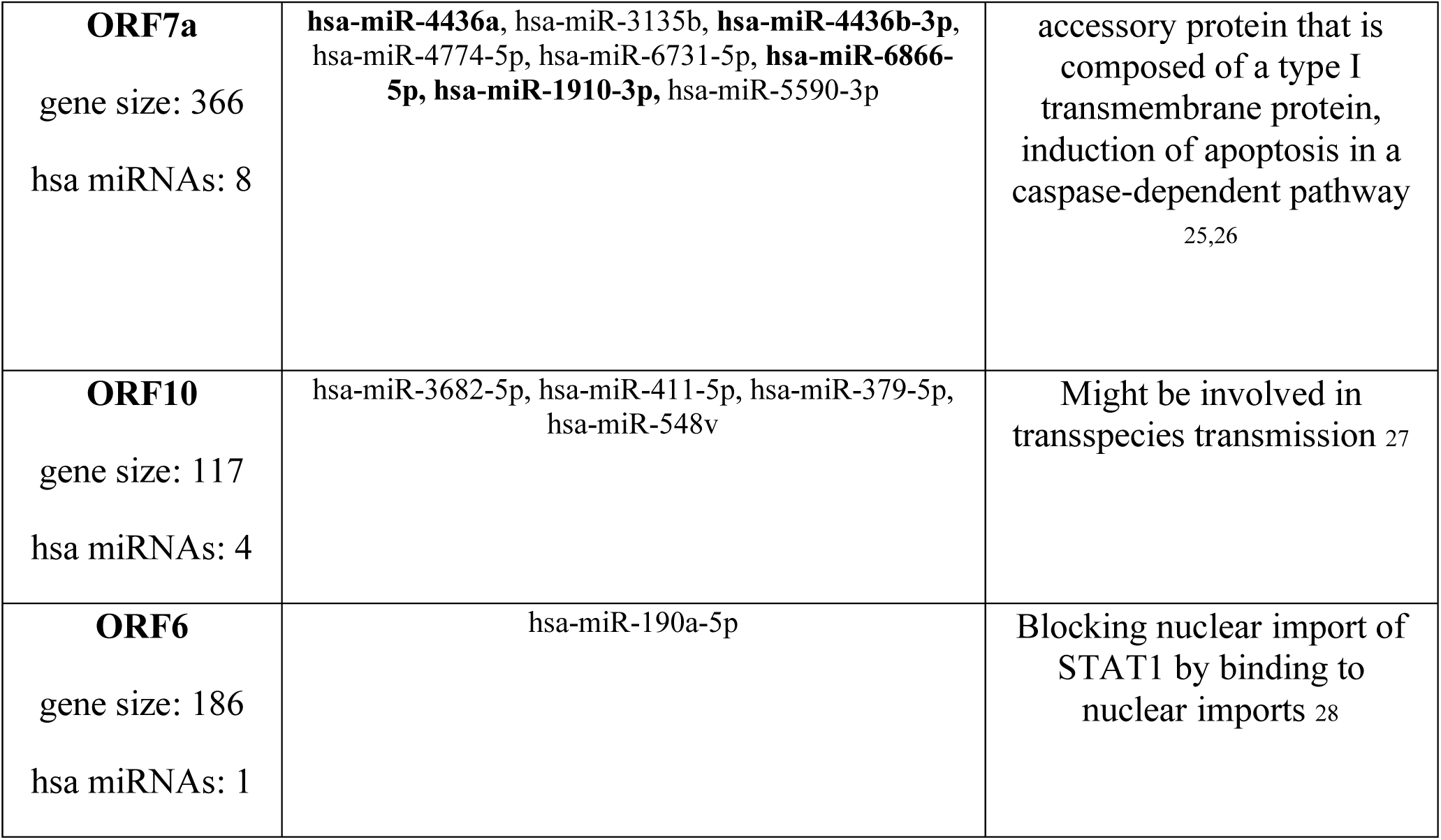
Predicted virulent mRNA targets by Human miRNAs: bold miRNAs are the common ones targeting more than one indicated viral proteins. The functions of SARS-CoV-2 proteins are not fully characterized, however, its coding genes might share functional similarity with SARS-CoV as shown in column “Functions of Target Genes”. Gene size indicates the size of genes in terms of number of nucleotides, hsa miRNAs shows the number of different human miRNAs that could target indicated viral genes.

Lastly, in order to understand the main mechanisms that would be affected by the influence of viral miRNAs on human genes, PANTHER Classification System was applied to targeted human genes. Based on the results presented in Figure 2 – 3 and Supplementary Files, a wide range of human genes with various molecular functions and pathways could be targeted.

**Figure 2:**
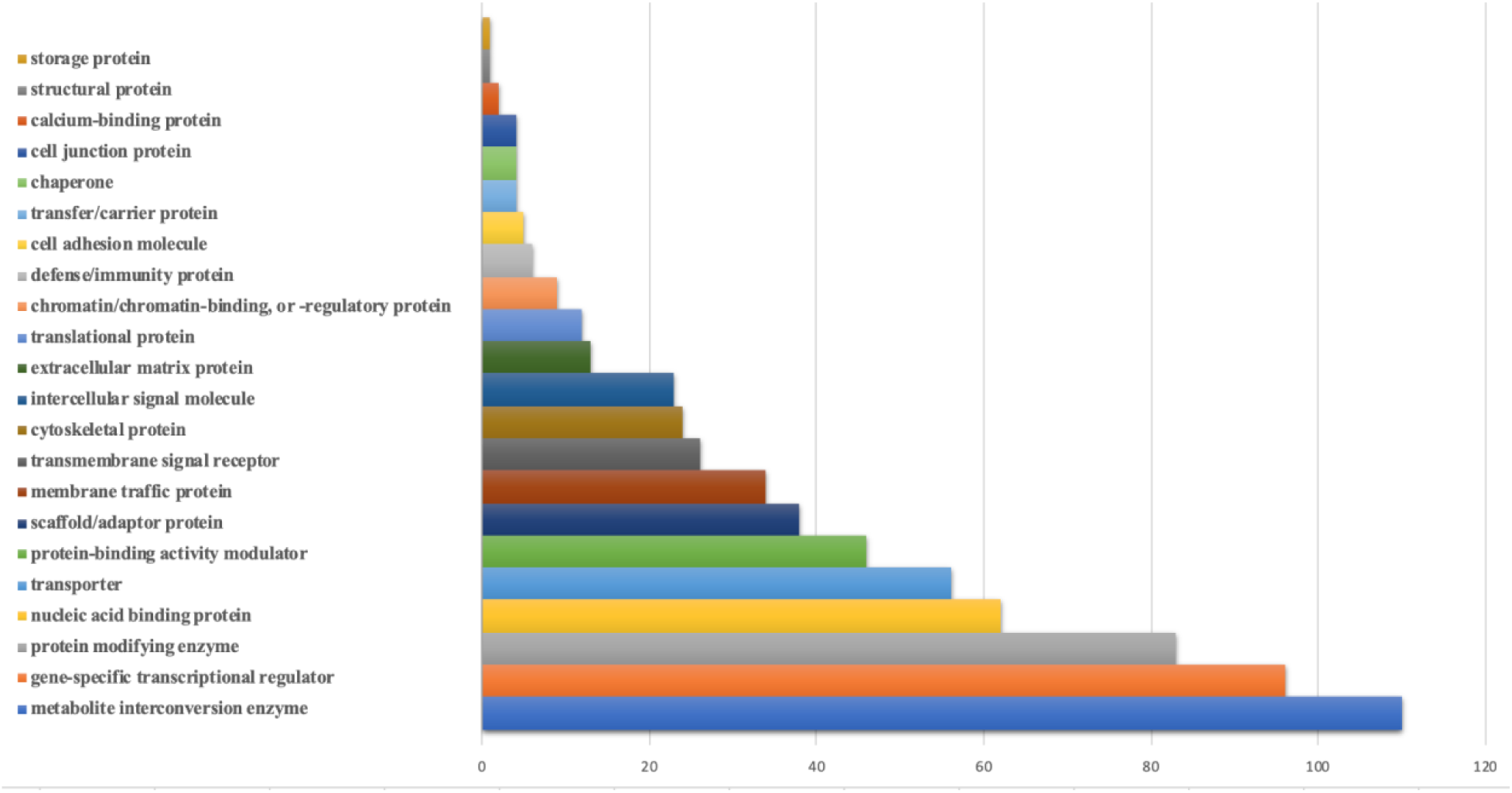
Bar-chart for the protein classes of human genes that could be targeted by viral miRNAs. Protein classes of genes were obtained from Panther. X-axis shows the number of genes with respected classes.

**Figure 3:**
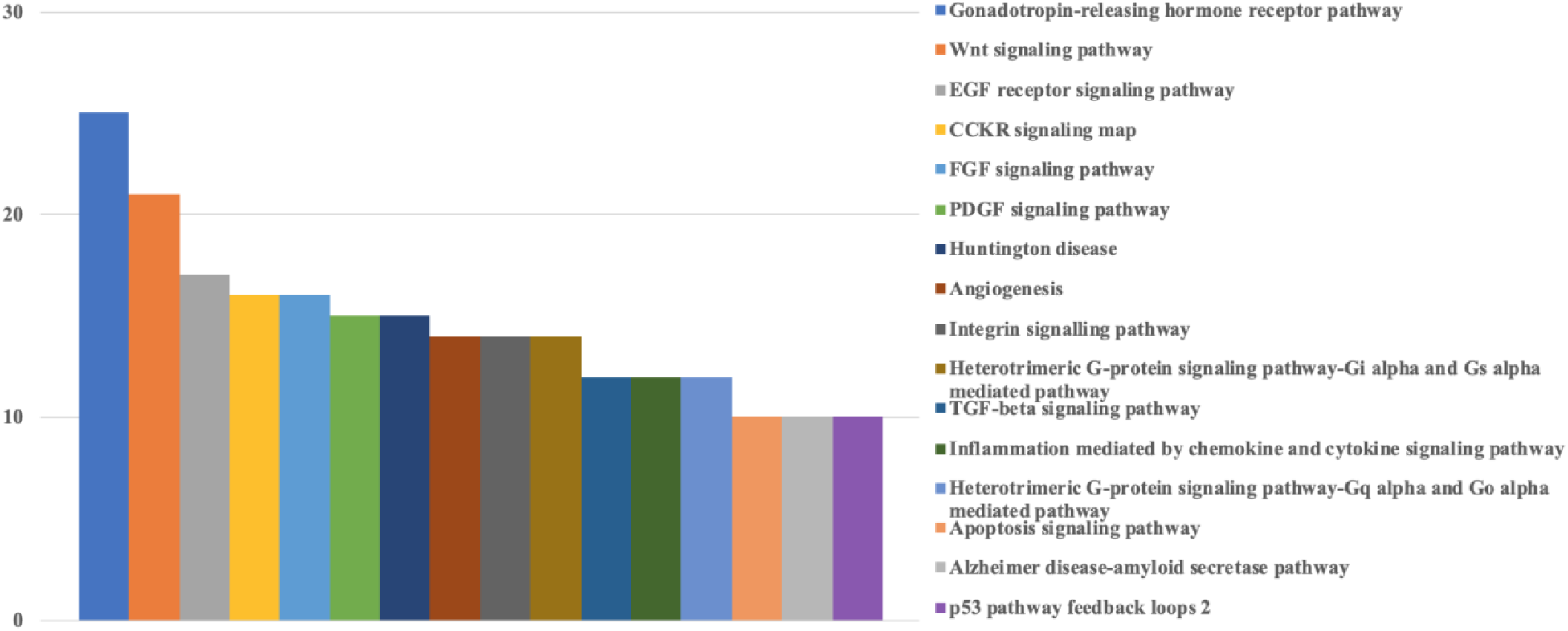
Bar-chart for the pathways of human genes that could be targeted by viral miRNAs. Graph is limited to the pathways that have at least 10 genes. Pathways of genes were obtained from Panther. Y-axis shows the number of genes with respected pathways. Legend is sorted from maximum to minimum (top to bottom).

### Discussion

The potential roles of miRNA-mediated RNA interference in infection biology has been defined as an essential regulatory molecular pathway. The involvement of miRNAs in host-pathogen interactions during infection might include the targeting of viral genes by host-cellular miRNAs as well as evolution of DNA and RNA virus-based gene silencing mechanisms to overcome host antiviral response and to maintain viral infection and disease ^29,30^. In addition, viruses might use host cellular miRNAs for their own advantage. Therefore, molecular elucidation of the roles of miRNAs in host-virus interaction might provide a deeper understanding for viral pathogenesis and the development of an effective antiviral therapy as shown in several viruses including Herpesvirus, Enterovirus and Hepatitis C ^30–32^. Although the detailed knowledge about viral miRNAs have been obtained from DNA viruses, it is still controversial for RNA viruses that whether they produce their own miRNAs or not ^33^. However, there are several reports showing the presence of non-canonical (due to the lack of classical stem-loop structure in miRNAs), small miRNA-like small RNAs produced during viral infections as shown in H5N1 Influenza ^34^, Ebola virus ^35^ and HIV-1 ^36^. Although they play crucial roles in several viral infections, miRNAs have not been studied in the pathogenesis of SARS-CoV-2 infection. In this study, we elucidated miRNA-mediated regulation responsible for SAR-CoV-2 infection by predicting possible host genes targeted by viral miRNAs and viral genes targeted by cellular miRNAs, which might provide potential ways to understand the underlying mechanisms of SARS-CoV-2 infection.

We investigated possible virulent genes of SARS-CoV-2 targeted by host-cellular miRNAs as shown in **Table 2**, which are mainly responsible for viral biogenesis, entrance, replication and infection. Except envelope (E) protein and ORF6, all viral genes (S, M, N, ORF1ab, ORF3a, ORF8, ORF7a and ORF10) are targeted by multiple human miRNAs. For instance, hsa-miR-203b-3p targeted ORF1ab and ORF3a with roles in viral replication was already shown to suppress influenza A virus replication ^37^. Even though hsa-miR-148a-3p targeted ORF8 to prevent interspecies transmission and also replication, it was found to target S, E, M and ORF1a protein in closely related SARS-CoV ^38^. hsa-let-7c-5p targeted ORF1ab in SARS-CoV-2 while it was found to be involved in H1N1 influenza A suppression by targeting its M1 protein ^39^. In another study, ORF6 of SARS-CoV suppressed type I interferon signaling by blocking the nuclear transport of STAT1 in the presence of interferon β ^40^, therefore, hsa-miR-190a-5p might target ORF6 to overcome immune system escape in SARS-CoV-2. The presence of such miRNAs could be considered as a host’s innate antiviral defense mechanism. On the other hand, virus could use these miRNAs to suppress their own replication to escape from immune system at the beginning of infection and transmission for a stronger infection. For instance, miR-146a-5p was upregulated in hepatitis C virus-infected liver cells of patients and in infected human hepatocytes, which promoted virus particle assembly ^41^. Moreover, ss RNA viruses could evolve very rapidly to change their gene sequences matching with these host miRNAs, therefore, they increase their host specificity. Once the virus establishes a successful transmission inside the host, they would mutate their genes very rapidly to escape from host miRNAs, which results from their RNA polymerases without proofreading activity ^42,43^.

Virus-derived miRNAs might function by targeting host and virus-encoded transcripts to regulate host-pathogen interaction. The roles of viral miRNAs in pathogenesis include alteration of host defense mechanisms and regulation of crucial biological processes including cell survival, proliferation, modulation of viral life-cycle phase ^44^. Although encoding miRNAs seems quite problematic for RNA viruses due to the nature of miRNA biogenesis pathway, it is possible to circumvent these problems through different ways as seen in HIV-1 ^36^. Therefore, we analyzed possible human genes targeted by predicted miRNA like small RNAs (**Table 1**). Based on the panther analysis, regulators of eukaryotic transcription would be the most important targets of 18 SARS-CoV-2 derived mature miRNA like candidates out of 29 hairpins in total. These transcriptional regulators are involved in both basal transcription machinery including several components of human mediator complex (MED1, MED9, MED12L, MED19) and basal transcription factors such as TAF4, TAF5 and TAF7L. Viruses might downregulate host gene expression in order to increase their gene expression either co-transcriptionally in the nucleus or post-transcriptionally in the nucleus or cytoplasm ^45^. Therefore, targeting basal transcription machinery such as components (TAFs) of TFIID complex by SARS-CoV-2 could prevent RNA polymerase II to assemble on promoters of host genes at the initiation step. Viral factors have been shown to block transcription initiation by inhibiting TFIID or more specifically TAF4 in herpesvirus ^46^. Another interesting target gene by SARS-CoV-2 miRNA was different subunits of CCR4-NOT transcription complex including CNOT4, CNOT10 and CNOT6L, which are deadenylases involved in mRNA decay ^47^. Therefore, suppression of these genes by viral miRNAs could impede mRNA turnover in the host and provide opportunities for viral mRNA to escape from degradation. Additionally, site specific trans-acting factors such as MAFG, STAT1, STAT5B and SOX11 would be targeted by viral miRNAs. STAT family transcription factors are activated by cytokine-induced stimuli and generally involved in an interferon response. In Kaposi’s sarcoma-associated herpesvirus (KSHV), viral miRNAs were found to inhibit STAT3 and STAT5, resulting in deregulated interferon response and transition into lytic viral replication ^48^.

As a conclusion, viral diseases have been paid attention as a global health problem due to the lack of proper treatment strategies and rapid evolution of viruses. In recent years, studies have focused on identifying miRNAs as targets for the treatment of viral diseases and there are promising clinical phase studies evaluating the therapeutic potential of antagomirs targeting host miRNAs such as miR-122 in hepatitis C ^49^. Based on the results shown in **Table 2**, it can be concluded that increases in the level of host miRNAs targeting virulent genes such as S, M, N, E and ORF1ab would block viral entry and replication. Moreover, decreasing the levels of host miRNAs would make SARS-CoV-2 more replicative and visible for the host immune system. However, alterations in host miRNA levels would interfere with specific cellular processes which are crucial for the host biology. In our study, we have also identified possible miRNA like small RNAs from SARS-CoV-2 genome which target important human genes. Therefore, antagomirs targeting viral miRNAs could be also designed even though there are only a few studies for DNA viruses ^45^. However, all these therapeutic possibilities need further mechanistical evaluations to understand how they regulate virus-host interaction. Therefore, further in vitro, ex vivo and in vivo studies will be required to validate candidate miRNAs for SARS-CoV-2 infection.

